# Low-temperature-specific effects of *PHYTOCHROME C* on the circadian clock in Arabidopsis suggest that *PHYC* underlies natural variation in biological timing

**DOI:** 10.1101/030577

**Authors:** Kieron D. Edwards, François Guerineau, Paul F. Devlin, Andrew J. Millar

## Abstract

The circadian clock is a fundamental feature of gene regulation and cell physiology in eukaryotes and some prokaryotes, and an exemplar gene regulatory network in Systems Biology. The circadian system in *Arabidopsis thaliana* is complex in part due to its photo-transduction pathways. Analysis of natural genetic variation between Arabidopsis accessions Cape Verde Islands (Cvi-0) and Landsberg erecta (L*er*) identified a major, temperature-specific Quantitative Trait Locus (QTL) on chromosome V that altered the circadian period of leaf movement (Edwards et al., Genetics, 2005). We tested Near-Isogenic Lines (NILs) to confirm that L*er* alleles at this *PerCv5c* QTL lengthened the circadian period at 12°C, with little effect at higher temperatures. The *PHYTOCHROME C* gene lies within the QTL interval, and contains multiple sequence variants. Plants carrying either a T-DNA-insertion into *PHYC* or a deletion of *PHYC* also lengthened circadian period under white light, except at 27°C. *phyB* and *phyABE* mutants lengthened period only at 12°C. These results extend recent data showing PhyC effects in red light, confirming the number of photoreceptor proteins implicated in the plant circadian system at eleven. The connection between light input mechanisms and temperature effects on the clock is reinforced. Natural genetic variation within *PHYC* is likely to underlie the *PerCv5c* QTL. Our results suggest that functional variation within the *PHYC-*L*er* haplotype group might contribute to the evolution of the circadian system and possibly to clock-related phenotypes such as flowering time. These results have previously passed peer-review, so we provide them in this citable preprint.

## INTRODUCTION

The circadian clock is a 24h endogenous timer that allows the correct temporal regulation of physiological, biochemical and developmental processes. The expression of over 30% of the Arabidopsis thaliana transcriptome is driven by the circadian clock (Covington et al. 2008; Michael et al. 2008; Michael and McClung 2003). Rhythmic transcriptional outputs control physiological processes such as daily rhythmic growth and photoperiodic flowering. In both cases, the mechanisms are sufficiently characterised to support mechanistic, mathematical models (Seaton et al. 2015). The clock mechanism in all organisms includes interlocked transcriptional–translational feedback loops. The clock’s rhythmic behaviour is thought to emerge from the dynamics of this clock gene circuit, which have been well characterised in Arabidopsis (Flis et al. 2015). The negative feedback loops in this model plant species incorporate two closely-related MYB transcription factors CIRCADIAN CLOCK ASSOCIATED1 (CCA1) and LONG ELONGATED HYPOCOTYL (LHY) that inhibit the expression of evening-expressed genes such as a pseudo-response regulator (PRR) TIMING OF CAB EXPRESSION 1 (TOC1). The expression of CCA1 and LHY is tightly regulated by other clock components, including sequential inhibition by PRR9, PRR7 and PRR5. TOC1 and other PRR genes are repressed by an Evening Complex (Hsu and Harmer 2014). For the clock to be useful, the endogenous period must be synchronised (entrained) to match the 24-hour environmental cycle (Johnson et al. 2003). The strongest entrainment signals are temperature and light. At least four families of photoreceptors have been identified as transducing light signals to reset the clock, the blue light sensing cryptochromes (*CRY1* and *CRY2*), the red/far-red light (R/FR) sensing phytochromes (*PHYA, PHYB, PHYD,* PHYE), (Devlin and Kay 2000; Somers et al. 1998; Yanovsky et al. 2000), the UV-B photoreceptor UVR8 (Feher et al. 2011) and a family of three F-box proteins, including *ZEITLUPE* (ZTL)(Baudry et al. 2010). These ten photoreceptors transduce light signals to regulate clock genes and proteins (Fankhauser and Staiger 2002), with both specialised and overlapping roles. *PHYC* alone has little effect on other phenotypes in the absence of other phytochromes (Hu et al. 2013), and no role for PhyC in the Arabidopsis clock had been confirmed until, during the preparation of this paper, a long-period phenotype was reported under red light (Jones et al. 2015). In barley, some *PHYC* alleles have also shown circadian effects (Pankin et al. 2014; Nishida et al. 2013) and in Arabidopsis, natural genetic variation in related traits has been associated with *PHYC* (Balasubramanian et al. 2006).

Temperature effects on circadian clocks include entrainment by temperature cycles, whereas under constant temperatures, the circadian period is unusually constant across a physiological temperature range (termed ‘temperature compensation’). Among multiple mechanisms implicated in how ambient thermocycles influence the clock are alternative RNA splicing, in *Neurospora crassa*, *Drosophila melanogaster* (Colot et al. 2005); Low *et al.* 2008) and Arabidopsis (James et al. 2012), and protein phosphorylation in *N. crassa* (Mehra et al. 2009). Mutants of some clock genes alter temperature compensation (Gould et al. 2006; Salome et al. 2010). A systems biology analysis of ambient temperature effects across the clock mechanism revealed a strong dependence on light quality, and suggested that light and temperature signalling converged upstream of the clock, such that photoreceptor pathways had a significant role in temperature responses (Gould et al. 2013). This work indicated multiple targets of temperature effects in the clock mechanism, including the morning genes implicated by Salome et al. 2010 and James et al. 2012, as well as the Evening Complex implicated by Mizuno et al. (2014).

Quantitative genetic approaches have identified genetic variation in clock-affecting genes (Anwer and Davis 2013), including Quantitative Trait Loci (QTL) in Arabidopsis (de Montaigu et al. 2015; Michael et al. 2003; Swarup et al. 1999). Our earlier work tested circadian period at three temperatures, identifying multiple QTL (Edwards et al. 2005). Here we propose *PHYC* as a candidate gene for a clock-affecting QTL that is specific for low ambient temperature.

### NILs recapitulate the temperature-specific QTL effect

The *PerCv5c* QTL was identified in the *Cvi* by L*er* (CvL) recombinant inbred lines (RILs) and mapped to the middle of Chromosome 5. This QTL alone accounted for 44.6% of phenotypic variation in the period of rhythmic leaf movement at 12°C (Edwards *et al.* 2005), by far the largest single effect observed in that study. Near isogenic lines (NILs) carrying Cape Verde Islands (Cvi) alleles around in the putative circadian QTL were therefore used to identify Cvi loci that regulate the clock, in an isogenic Landsberg *erecta* (L*er*) background (Edwards et al. 2005). Figure 1a shows the periods of NIL45a and NIL106 at 12°C, 22°C and 27°C, with numerical values in Table 1. Both NILs had Cvi introgressions around the *PerCv5c* locus (Figure 1b). Both NILs had shorter circadian period compared to the L*er* parent especially at 12°C (Figure 1a), consistent with the original QTL identification. The larger introgression of NIL45a shortened the period more than NIL106. It is possible that multiple loci in the Cvi region of NIL45a affect the clock but the small period effects make this difficult to confirm.

**Table 1.**
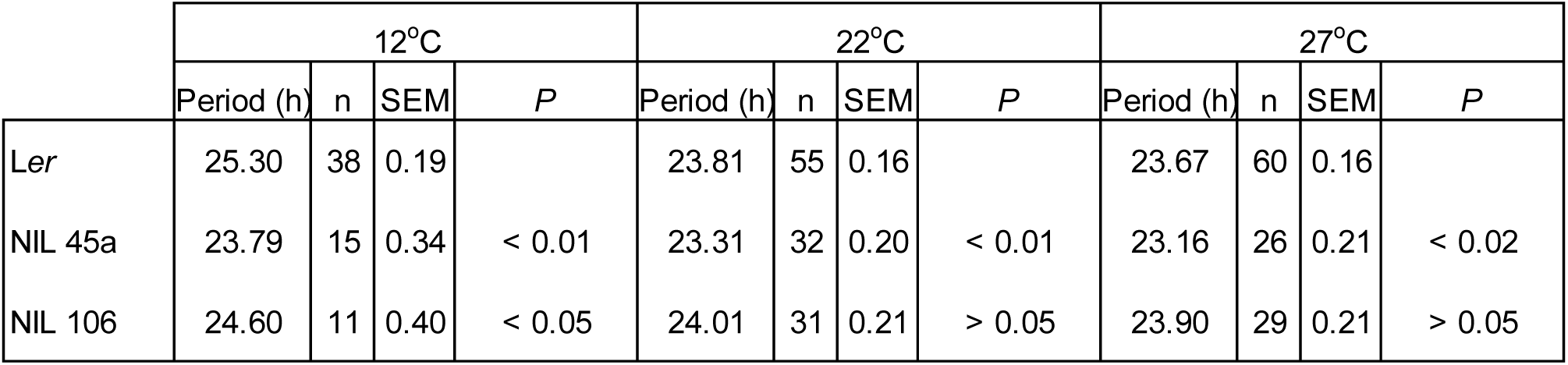
Summary of *PerCv5c* NIL leaf movement periods. Leaf movement period of *PerCv5c* NILs at 12°C, 22°C and 27°C indicated. Data are means of n traces per line at each temperature. Significance levels of *t*-tests comparing the mean periods of NILs to L*er* are shown (*P*).

|  | 12°C |  |  |  | 22°C |  |  |  | 27°C |  |  |  |
| --- | --- | --- | --- | --- | --- | --- | --- | --- | --- | --- | --- | --- |
|  | Period (h) | n | SEM | *P* | Period (h) | n | SEM | *P* | Period (h) | n | SEM | *P* |
| *Ler* | 25.30 | 38 | 0.19 |  | 23.81 | 55 | 0.16 |  | 23.67 | 60 | 0.16 |  |
| NIL 45a | 23.79 | 15 | 0.34 | < 0.01 | 23.31 | 32 | 0.20 | < 0.01 | 23.16 | 26 | 0.21 | < 0.02 |
| NIL 106 | 24.60 | 11 | 0.40 | < 0.05 | 24.01 | 31 | 0.21 | > 0.05 | 23.90 | 29 | 0.21 | > 0.05 |

**Figure 1.**
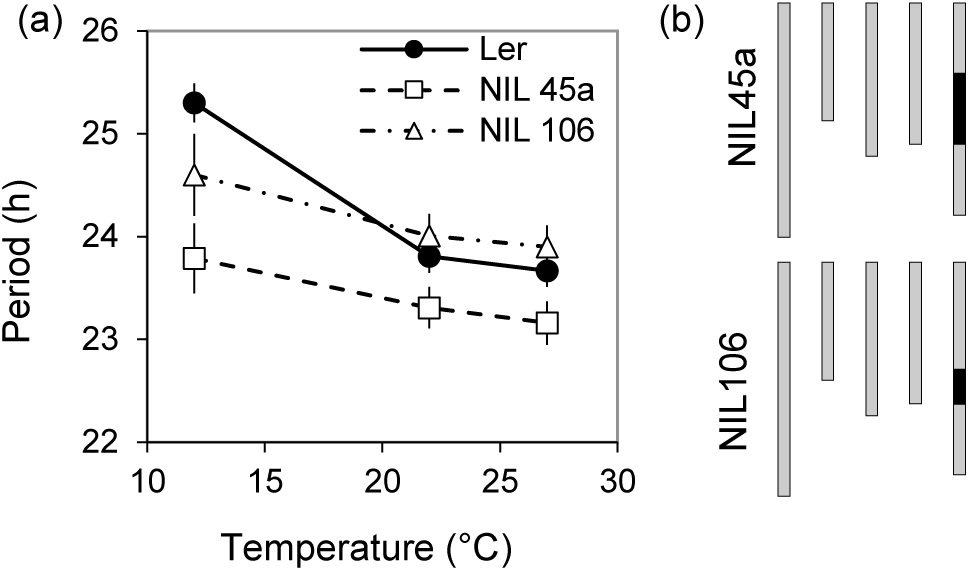
*PerCv5c* QTL effect on period, defined in Near-Isogenic Lines (NILs). (a) *PerCv5c* NIL periods. Mean leaf movement period versus temperature of *PerCv5c* NILs 45a (open squares) and 106 (open triangles) compared to L*er* (filled circles) at 12°C, 22°C and 27°C. Error bars represent SEM of period estimates. (b) Graphical genotypes of the NILs. Vertical bars represent linkage groups and colour represents L*er* (grey) background or Cvi (black) genome sequences on chromosome V.

### Natural allelic variation at *PHYTOCHROME C*

*PHYC* maps within the confidence interval of *PerCv5c,* so was considered a possible candidate gene for the QTL. Quantitative RT-PCR showed that the *PHYC* alleles in both accessions were expressed under constant light conditions (Figure 2). Expression was higher in subjective day (54h in constant light) than in subjective night (62h in constant light), though possibly with lower levels of mRNA in *Ler* (Figure 2).

**Figure 2.**
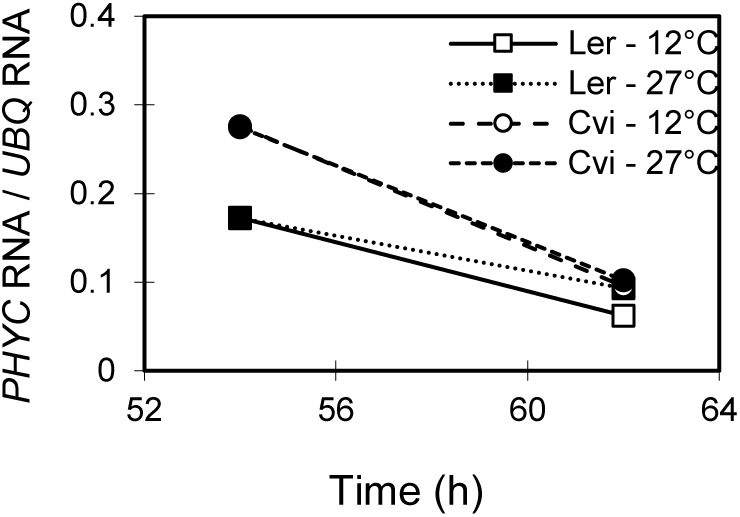
*PHYC* expression in Cvi and L*er*. Quantitative RT-PCR showed that both alleles were expressed, in RNA extracted from Cvi and from L*er* plants at the times (54h, 62h) and temperatures (12°C, 27°C) indicated, though possibly with lower levels of mRNA in L*er*.

Sequence analyses for the *PHYC* gene and predicted PHYC protein revealed mutations between the *Cvi* and L*er* backgrounds, leading to four changes in the protein sequence (Figure 3a, 3b). The AT5G35840 locus in sequences from the 1001 Genomes Project (Weigel and Mott 2009) confirmed these four changes (Figure 3c). The 1001 Genomes resource also showed that the Cvi sequence is close to the L*er* haplotype rather than the Col reference haplotype, which is shared by Ws-2. The changes from L*er* to Cvi amino acid sequence are A27T, S230G, T352S and H470P, all of which fall in the N-terminal domains of phytochrome before the bipartite PAS motif. At position 230, G is the amino acid in the reference Col genome, whereas in the other positions, the Col sequence is the same as L*er* (Figure 3c).

**Figure 3.**
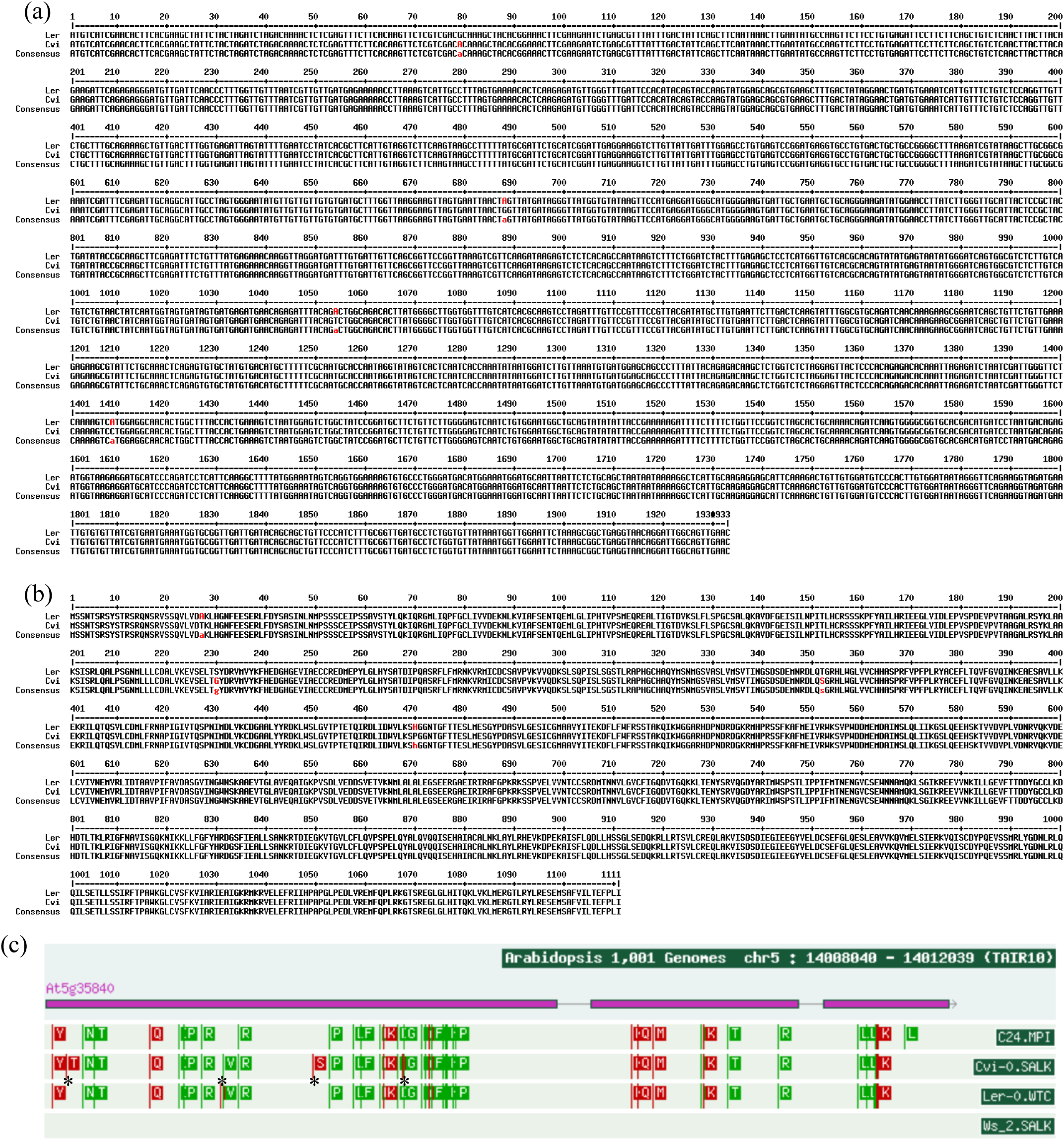
Genomic and protein sequence alignment in Cvi and L*er.* Sequence analyses from the plants tested are shown for the (a) *PHYC* gene and (b) predicted PHYC protein, revealing variation between the *Cvi* and *Ler* backgrounds leading to four amino acid changes in the protein sequence (L*er* -> Cvi): A27T, S230G, T352S and H470P. For S230G, the L*er* amino acid differs from the Col-0 reference. (c) corresponding analysis of *PHYC* locus AT5G35840 from the 1001 Genomes Project server (http://signal.salk.edu/atg1001/3.0/gebrowser.php), showing the four accessions relevant to this study: Cvi-0 and L*er* from the CvL RILs and NILs, and the C24 and Ws-2 genetic backgrounds for the *phyc* mutant alleles. Asterisks between Cvi-0 and L*er* mark the four amino acid substitutions detected in (b). Ws-2 is identical to the Col-0 reference sequence. Note the additional L substitution at the C terminus of the C24 allele. Red markers, A in reference sequence; Green markers, G in reference sequence. Alternative sequences are provided for Cvi-0 and L*er* in this resource (not shown) but they did not match our sequences in (a).

### *phyC* mutants alter circadian period

To test a possible role for *PHYC* in the circadian clock, the *phyC-1* mutant, in a PhyD-deficient background Wassilewskija-2 (Ws) (Franklin et al. 2003) was assayed for period at 12°C, 22°C and 27°C (Figure 4a). *phyC-1* displayed 1.6 and 1.4 hours period lengthening at 12°C and 22°C respectively, but was not significantly different from wild type at 27°C. Numerical period values and statistical tests are shown in Table 1.

**Figure 4.**
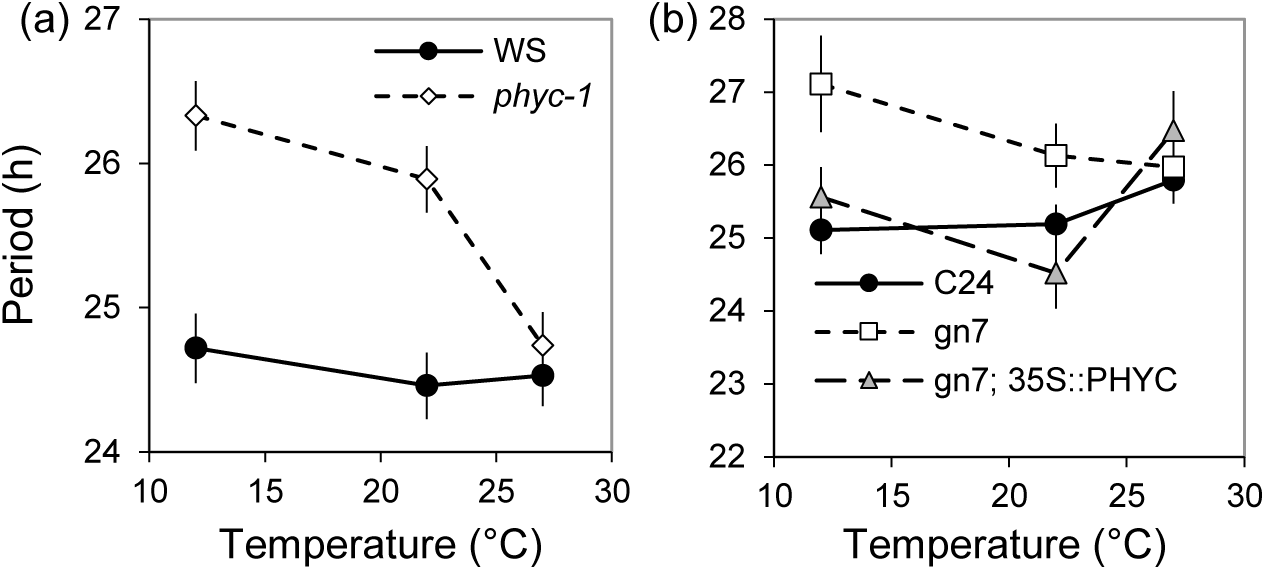
PhyC controls circadian period at low ambient temperature. (a) Leaf movement period of *phyC-1* mutant plants compared to wild type Ws. (b) Leaf movement period of C24 wild type (filled circles), gn7 deletion mutant (open squares) and gn7; *35S::PHYC* rescued line (open triangles) at 12°C, 22°C and 27°C. Error bars represent SEM of period estimates.

To confirm the *phyC-1* phenotype, period was tested in a second null mutant for *phyC.* The gn7 line, first described as gne7 by Sorensen *et al.* (Sorensen et al. 2002), has a large genomic deletion removing the entire *PHYC* open reading frame in a C24 background (Figure 5a). As with *phyC-1*, significant circadian period lengthening was observed in gn7 plants at 12°C and 22°C, but not 27°C (Figure 4b), relative to the C24 parent line. Over expression of *PHYC* under the 35S promoter rescued this period phenotype, suggesting that the gn7 period effect was principally mediated by the deletion of *PHYC* (Figure 4b).

**Figure 5.**
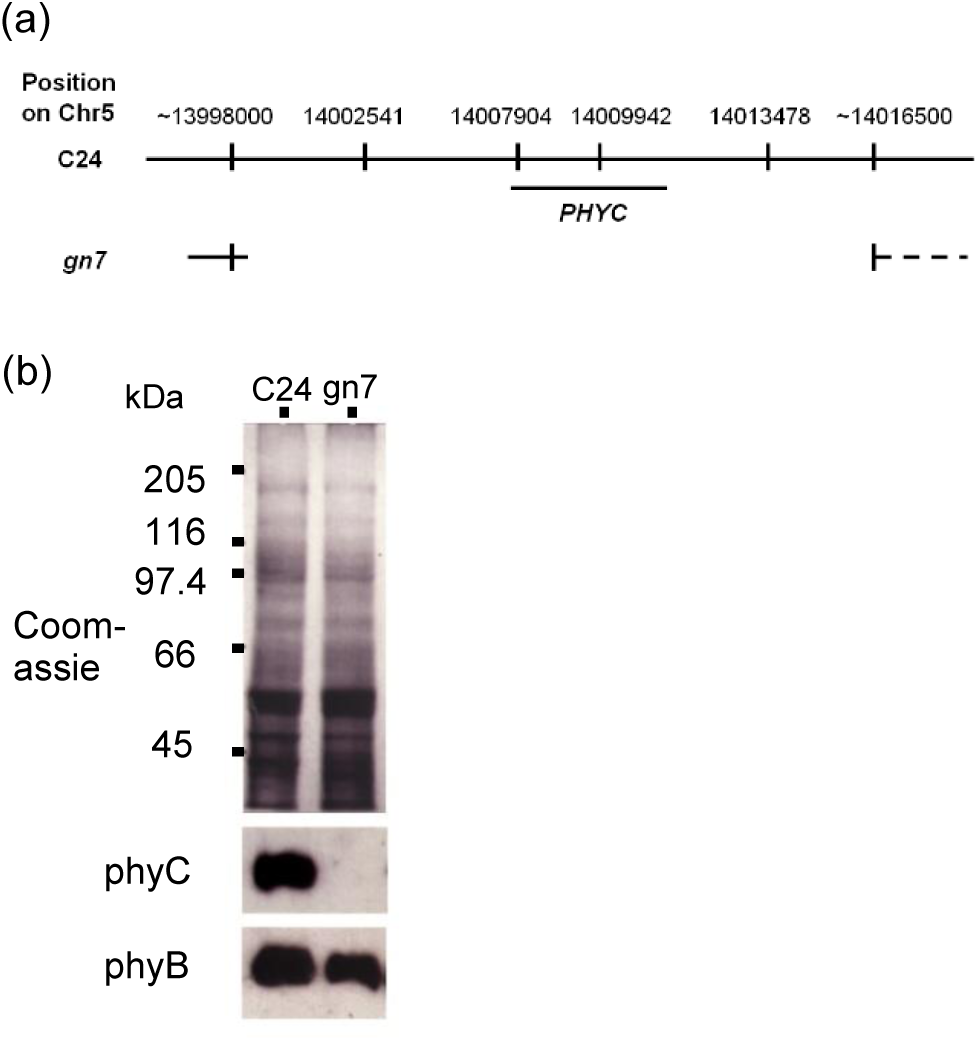
The *PHYC* open reading frame is completely removed in the gn7 line, (a) Schema of deletion in gn7 as detected by Southern probes at locations indicated. (b) Western blotting performed on protein extracts of C24 and gn7 plants, for phyC and phyB proteins confirms that phyC protein is specifically absent in *gn7.* The Coomassie-stained control confirms equal protein loading.

*phyB-10* single mutants and *phyABE* triple mutants were tested in the same experiments. Neither mutant showed a period defect in white light at 22°C, as expected, but both lengthened period at 12°C (Figure 6; Table 2).

**Figure 6.**
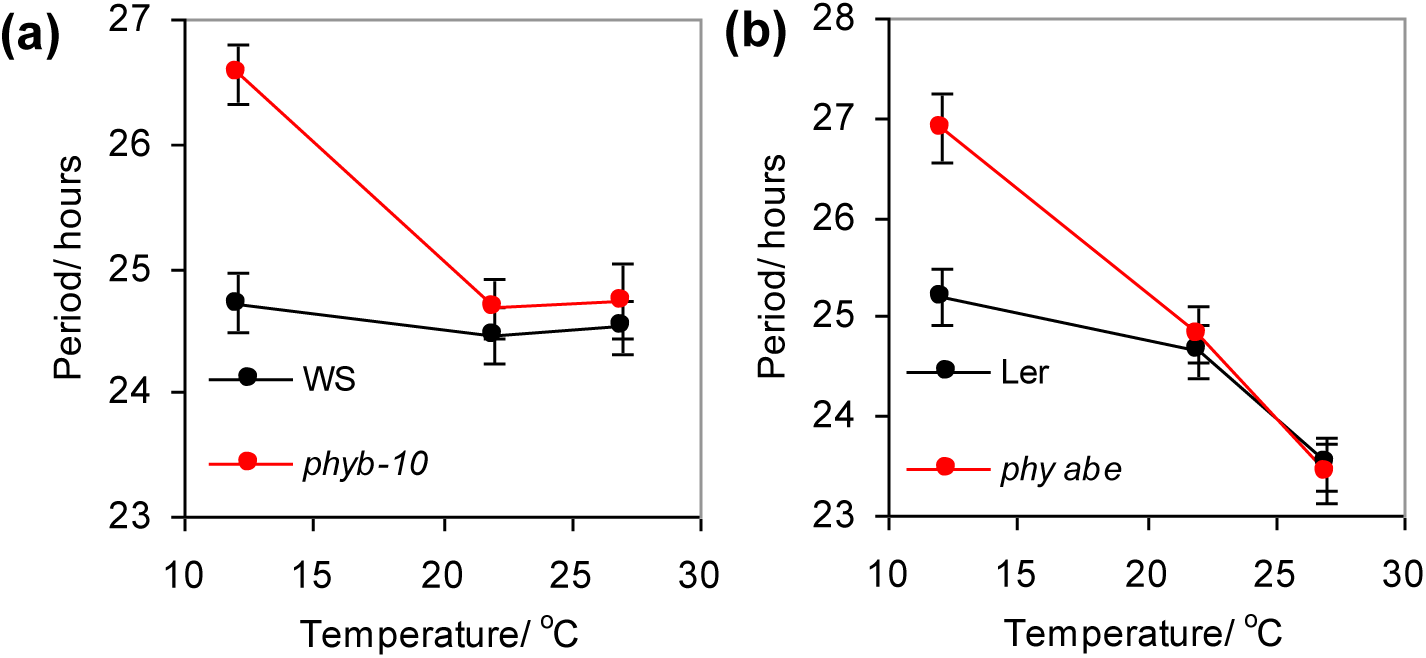
Circadian period of *phytochrome B* single and multiple mutants. Leaf movement period of (**a**), *phyb-10* (b) *phy abe* mutants compared to respective wild types. See inset legends for line identity. Error bars represent SEM of period estimates.

**Table 2.**
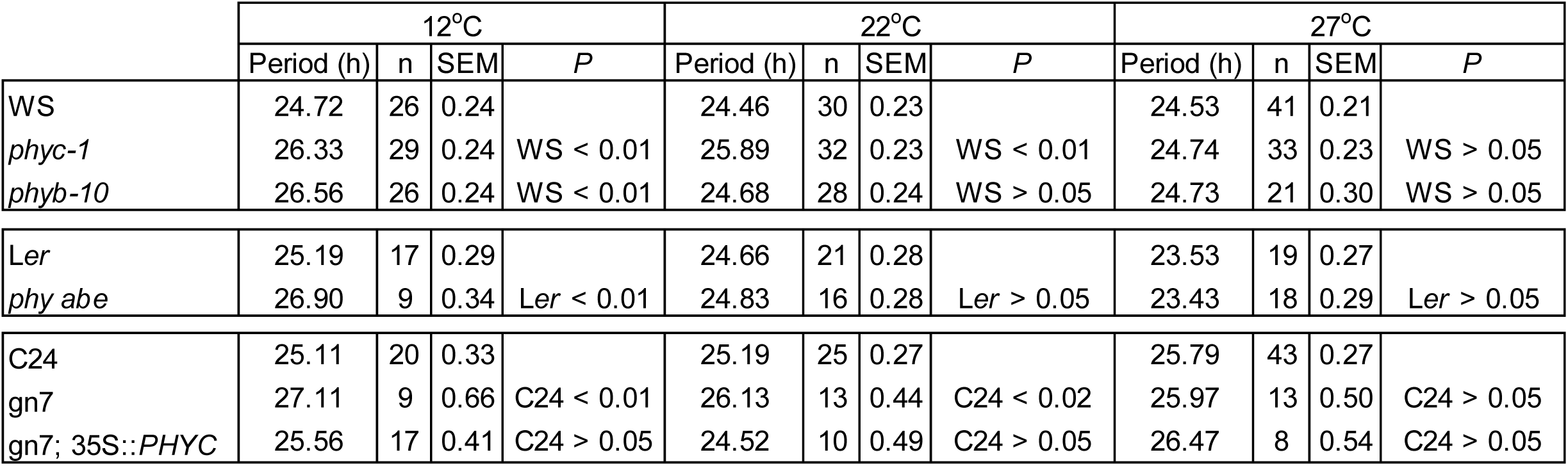
Summary of phytochrome mutant leaf movement periods. Leaf movement period of *phy* mutants at 12°C, 22°C and 27°C indicated. Data are means of n traces per line at each temperature. Significance levels of *t*-tests comparing the mean periods of mutants to wild type are shown *(P).*

## DISCUSSION

Circadian clock mechanisms include gene regulation by multiple, interlocking feedback loops, which can increase the flexibility of possible regulatory changes over evolutionary time and in the face of environmental variations. In light-grown Arabidopsis seedlings, multiple photoreceptors contribute light input signals, adding further complexity to the clock network.

We showed that phyC contributes to control clock period in a temperature-specific manner under white light, adding to the four other phytochromes, two cryptochromes, UVR8 and the ZTL/FKF1/LKP2 family, and bringing the number of known circadian photoreceptor proteins in Arabidopsis to eleven. The rhythmic control of native *PHYC* expression (Toth *et al.* 2001) was not required for normal circadian rhythmicity, because the gn7 mutant phenotype was rescued by a 35S mis-expression transgene (Figure 4b).

A similar phenotype was observed in the *phyb-10* and *phyabe* triple mutant, suggesting that this effect was principally mediated by *phyb* in both lines. In contrast to the *phyB* mutants, both *phyC* mutants lengthened period at 22°C as well as 12°C. Taken together, these phenotypes suggested that the *phyD* mutation in the Ws background of *phyc-1* and *phyb-10* was not uniquely required for the phenotypes observed, though interactions among the phytochromes might well contribute. Hence, *PHYB* and *PHYC* are likely the major effectors of the clock regulation detected here.

No circadian period QTL were mapped to the *PHYB* locus in Edwards et al. (2005). However, an epistatic interaction was suggested between markers FD.222L-Col and CH.60C, which map near to *PHYB* and *PHYC* respectively (K.D. Edwards Ph.D. Thesis, University of Warwick). This interaction suggested the possibility of the L*er* allele of *PHYB* enhancing the period difference between the L*er* and Cvi alleles of *PHYC.*

The similarity of the *phyC* mutant phenotype to the *PerCv5c* QTL effect, together with the DNA sequences, strongly suggested that allelic variation at *PHYC* contributes to or solely causes this QTL. If so, then our results further suggested that the *PHYC-*L*er* allele was less active than *PHYC-Cvi,* consistent with (Monte et al. 2003; Balasubramanian et al. 2006), because *PHYC-*L*er* lengthened circadian period as *phyC* null mutants also do. The haplotypes of *PHYC* across multiple accessions were defined by Balasubramanian et al. (2006). Cvi is clearly among the L*er* haplotypes, whereas Ws-2 is among the Col haplotypes (Figure 3c). Our results suggest that *PHYC* function varies within the L*er* haplotype group, at least at low temperature. The variation within this group may be relevant, for example, to explain the association of *PHYC* alleles within the L*er* haplotype group to particular habitats, such as areas of high precipitation in the Iberian peninsula (Mendez-Vigo et al. 2011).

Page 6, below, shows **Figure 6. Circadian period of *phyB* single and multiple mutants. Table 2. Summary of leaf movement periods in** *phy* **mutants of Figs. 4 and 6.**

## EXPERIMENTAL PROCEDURES

### Plant material

Seeds for Arabidopsis accessions, CvL RILs and NILs used in leaf movement analysis were as described (Edwards et al. 2005), generously donated by M. Koornneef. NIL45a carries Cvi alleles in a 42-48 cM region in the middle of Chromosome 5, between markers GH. 177C and HH.4451Col (Alonso-Blanco et al. 1998). NIL106 has a 15-19 cM Cvi introgression between markers GB.235-Col and GB.233C (Alonso-Blanco et al. 1998). *phyC-1* null allele contains a T-DNA insertion as previously described (Franklin et al. 2003). *phyC* deletion allele gn7 has previously described (Sorensen et al. 2002); seeds for gn7 and gn7 expressing *35S::PHYC* were generously provided by G. Whitelam. *phyB-10* mutants were previously named *phyB-464-19* (Reed et al. 1993) in the Ws background. The *phyabe* mutant contains the *phya-2* (Whitelam et al. 1993), *phyb-1* (Koornneef et al. 1980) and *phye* (Devlin et al. 1998) alleles in a L*er* background.

### Growth conditions

Growth conditions for samples used in leaf movement and RNA studies under constant light were essentially as described (Edwards et al. 2005; Edwards et al. 2010). Briefly, sterile seed were stored in 0.15% agar and stratified at 4° for 4–5 days prior to sowing on Murashige-Skoog 1.5% agar medium containing 3% sucrose. Seedlings were grown for 6 days under constant light of 55–60 μmol m−2 sec−1 cool white fluorescent light and then entrained for 4 days at 21°−22° under (12 hr/12 hr) light/dark cycles of 75 μmol m−2 sec−1 cool white fluorescent light.

### Measurement of leaf movement

Individual period estimates were produced from leaf-movement data as described (Edwards et al. 2005). In brief, leaf growth of Arabidopsis seedlings was imaged under constant light using a custom-built, 31-camera array. Y-axis centroid positions for each leaf were determined from the image stacks using Metamorph software. Circadian period of each leaf was estimated using the FFT-NLLS algorithm through the BRASS interface (Edwards et al., 2010). Mean period estimates for each genotype were based on data from two to four independent experiments at each temperature analysed using REML (Patterson and Thompson 1971) in the statistical package GENSTAT 5 (Payne et al. 1993) as described (Edwards et al. 2005). The significance of differences between pairs of genotypes was analysed via t-tests using the SEM estimates derived from REML.

### Molecular Assays

Quantitative RT-PCR was performed according to (Edwards et al. 2010). Phytochrome Western blotting was performed as described (Franklin et al. 2003).

Citation: K.D. Edwards, F. Guerineau, P.F. Devlin and A.J. Millar (2015) Low-temperature-specific effects of PHYTOCHROME C on the circadian clock in Arabidopsis suggest that PHYC underlies natural variation in biological timing. bioRxiv, DOI: 10.1101/030577

## Author contributions

KDE and AJM designed the study. FG characterised the gn7 deletion in the Scott lab (Figure 5a). PFD analysed PHY protein content in the gn7 line (Figure 5b). KDE performed all other experiments and data analysis. AJM assembled the data, removed them from Wenden et al. Plant Journal 2011 after peer review to comply with an editorial request for greater focus, and prepared this paper.

## Acknowledgements

We are grateful to Rod Scott and the late Garry Whitelam for supporting early work on gn7, and to Dr. James Lynne (Horticulture Research International, Wellesbourne) for REML analysis.

